# Fossil data do not support a long pre-Cretaceous history of flowering plants

**DOI:** 10.1101/2021.02.16.431478

**Authors:** Graham E. Budd, Richard P. Mann, James A. Doyle, Mario Coiro, Jason Hilton

## Abstract

The origin of angiosperms is a classic macroevolutionary problem, because of their rapid rise in the Early Cretaceous fossil record, beginning about 139 Ma ago, and the conflict this creates with older crown-group ages based on molecular clock dating^1^. Silvestro et al.^2^ use a novel methodology to model past angiosperm diversity based on a Bayesian Brownian Bridge model of fossil finds assigned to extant families, concluding that a Cretaceous origin is vanishingly unlikely. However, their results strongly conflict with the known temporal distribution of angiosperm fossils, and, while we agree that statistical analysis aids interpretation of the fossil record, here we show the conclusions of Silvestro et al.^2^ are unsound.

---

The outputs of the new method^2^ are strikingly well predicted simply by the age of oldest fossil and the number of fossils in each family (Figs 1A, 2), just as in the older stratigraphic confidence interval estimation method^3^, and plotting results of the two methods against each other reveals an extremely close correspondence (Fig. 3). In either method, estimates of the true range of a group based on fewer than six fossil occurrences are highly uncertain. Conversely, with six or more fossils, both methods converge to a central estimate close to the age of the oldest fossil. The novel components of the new method^2^, in particular the stochastic model of diversification, are thus largely redundant, and do not extract any more information from the data.

**Figure 1.**
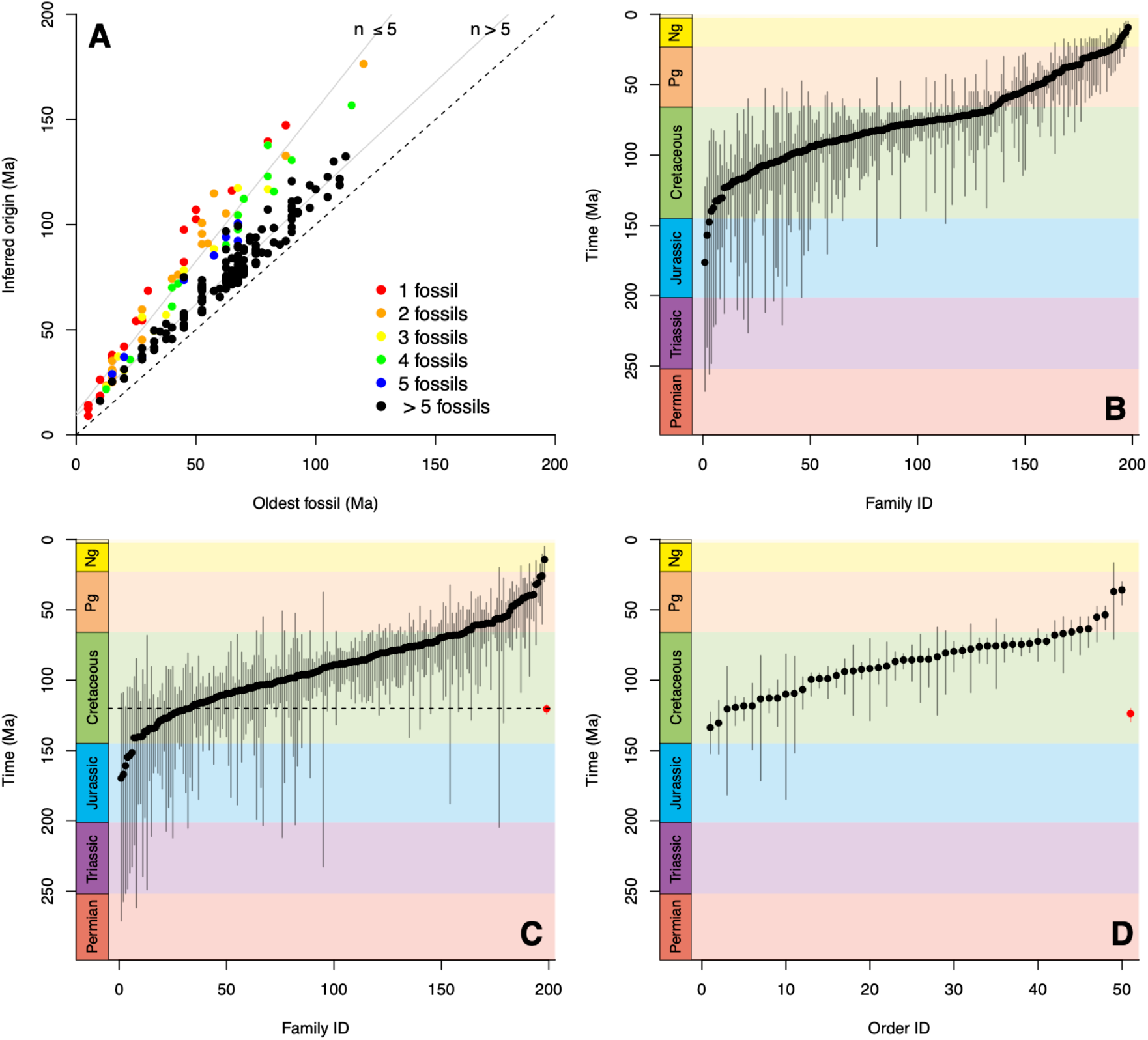
**A** Plot of mean inferred age for each of 198 families from the analysis of ref. 2 against age of oldest fossil for that family, colour-coded to show number of fossils in each family. Regression lines show the dependence of inferred age on number of fossils for 1-5 fossils (y = 10.64x + 1.44; r^2^ = 0.95) and more than 5 (y = 9.10x + 10.6; r^2^ = 0.93), showing that nearly all the variation is captured by these two variables. Note that regression on individual number of fossils typically increases r^2^ values even further (see Extended Data Fig. 2). **B** inferred means and 95% credible intervals of 198 angiosperms of the dataset and analysis of ref 2., with a 2.5 Myrs time bin and including their pollen records. The Jurassic-Cretaceous boundary is marked with a dashed line. **C** A similar analysis, but performed on a simulated data set of 198 “families” each created from a birth-death process of age 120 Ma (dashed line – the age of the oldest fossil in the analysis of ref. 2), with an extinction rate of 0.5 and speciation rate of 0.5349469 to create an expected 1000 species at the present day. Fossils were populated using a binomial sampling of the diversity through time at rate q = 0.001, giving on average 12 fossils per “family”, and 68 “families” with five or fewer fossils (note that this broadly matches the 72 families with five or fewer fossils in the analysis of ref. 2). Note the striking resemblance to panel **B. D** Analysis of ref 2. as in panel **B**, but performed at the order, not family level (aggregate plot in red). Note that all mean probabilities now lie within the Cretaceous (see also Extended Data Figs 1, 2). Panels **C, D** plotted using the code of ref. 2 with default settings, with q_var = 0 and 1 respectively.

**Figure 2.**
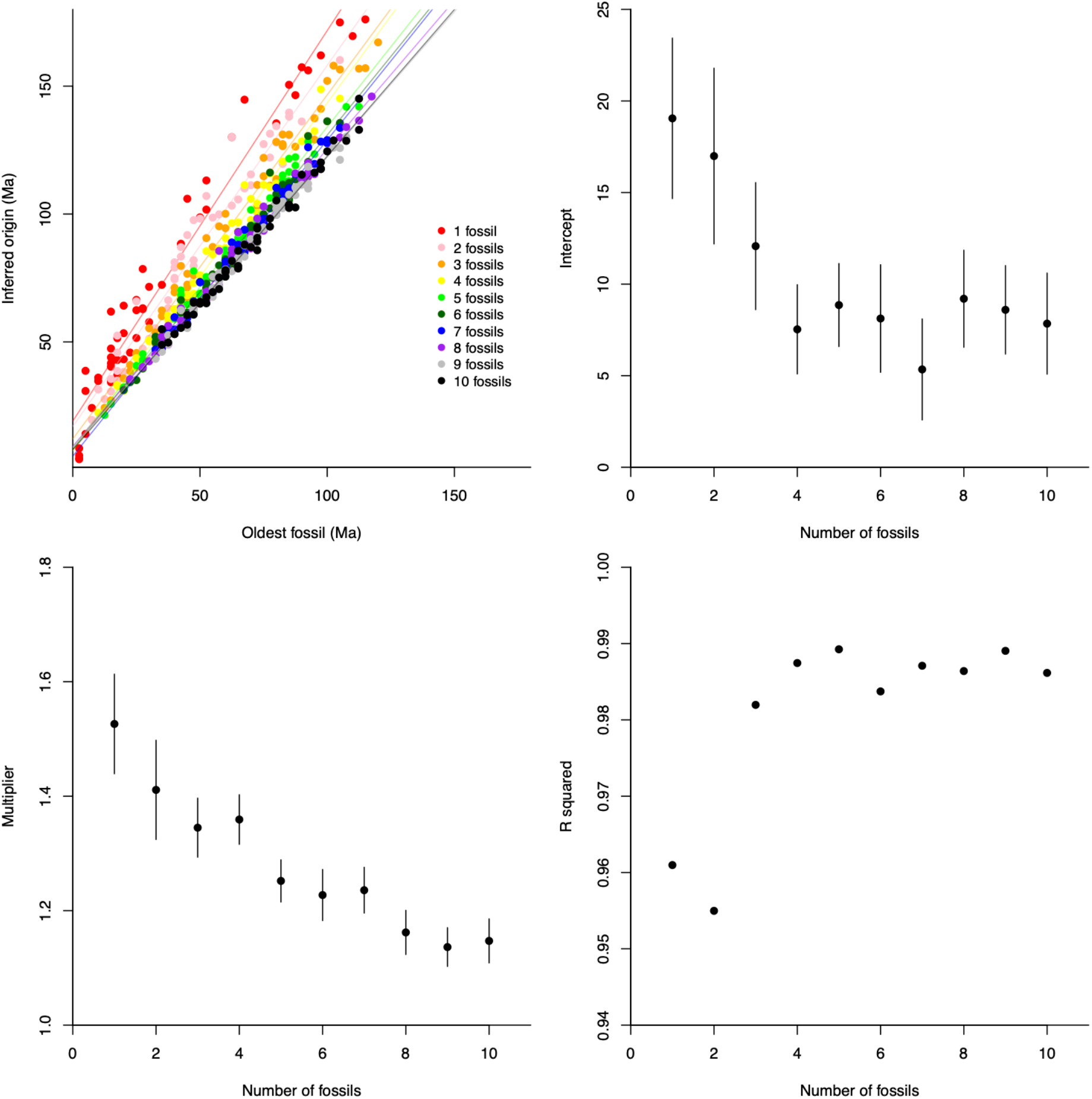
**A** Simulated sets of 20 families, selected to have 1, 2, 3 etc fossils showing regression lines of inferred age on age of oldest fossil. **B** Intercepts of the regression lines in **A** against number of fossils. **C** Slope of regression lines from **A** against number of fossils. **D** r^2^ values for the regression lines in **A**. Note that almost all the variation is explained only by age of oldest fossil and number of fossils.

**Figure 3.**
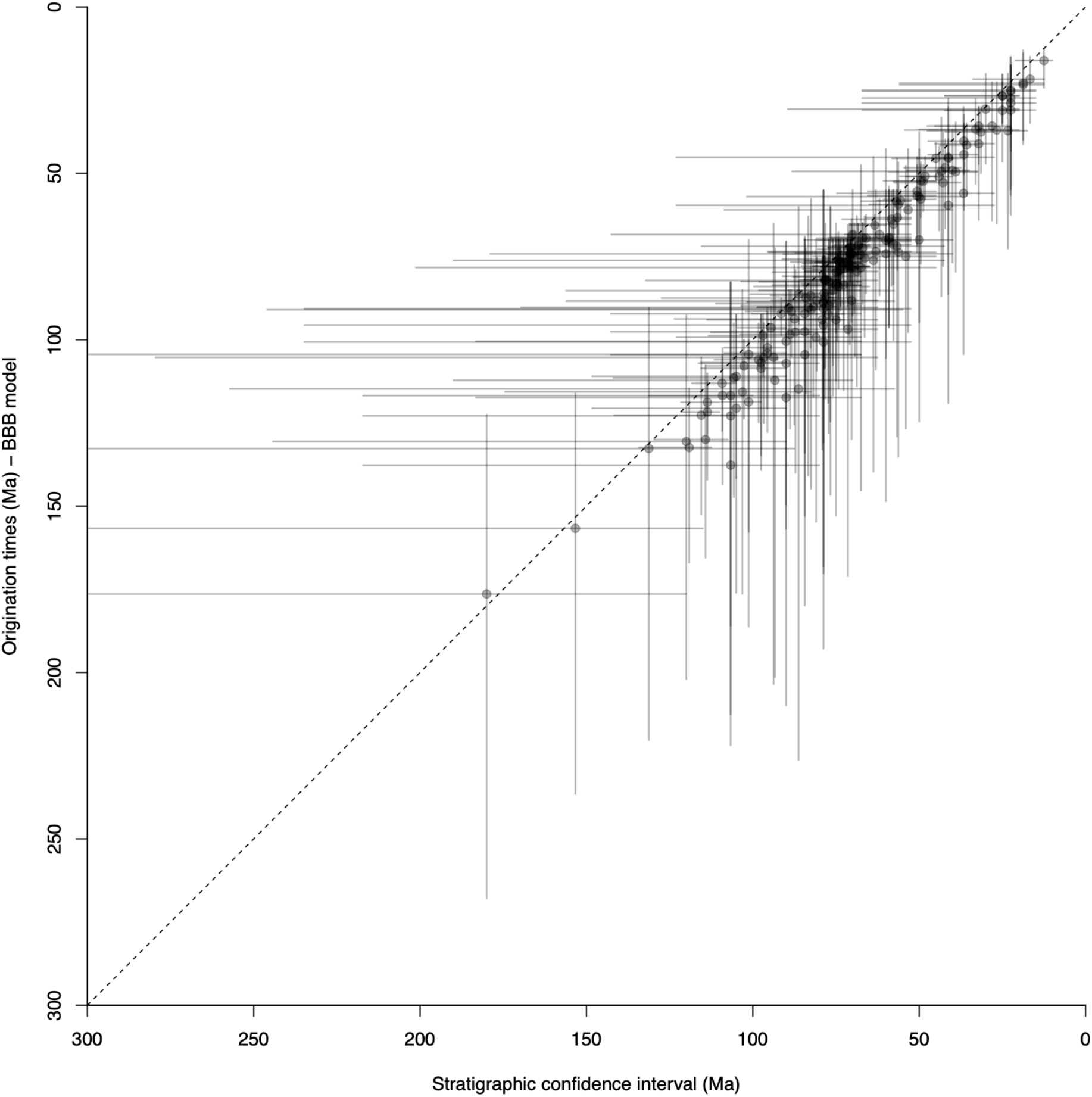
Origination times estimated by the BBB model^2^ compared to stratigraphic confidence estimates of ref. 3, using code adapted from ref. 2 (samples with only one fossil horizon are omitted as in fig. 5b of ref. 2). Note that we identified the following issues in the equivalent calculation performed in ref. 2 fig. 5b: (i) since these are extant families, the range (although not number) of observed instances for each family should extend to the present -ref. 2 instead takes the range as being limited by the most recent fossil; (ii) we plot instead of the midpoint of the confidence intervals (which have no special significance) the unbiased minimum variance estimator^3^. Note that with these corrections, the differences between the methods of ref. 2 and ref. 3 are minimized.

To test whether or not the results of ref. 2 support their conclusions, we simulated 198 clades (the number of angiosperm families in ref. 2) using a birth-death process^4^, each originating at 120 Ma (i.e. in the Cretaceous) and populated with fossils by binomial sampling from their true diversity through time. We then attempted to recover the time of origin of each simulated family using the methodology and code of ref. 2. This produced a pattern strikingly similar to that in ref. 2 (Fig. 1B, C) – despite our simulated families all being 120 Myrs old. Thus, their results are entirely compatible with an origin of angiosperms in the Cretaceous. In both cases, the apparent pattern of diversification can be ascribed to inherent variability in the age of the oldest fossil (even amongst families of the same age), which is greatly accentuated in the sparsely preserved families that dominate both the oldest and youngest inferred origins (compare Figs 4A and B). In short, the Jurassic ages of the oldest families inferred in ref. 2 are generated by their sparse records and the vagaries of stratigraphic sampling (these ones happening to have relatively deep oldest fossils), so their range is uncertain.

**Figure 4.**
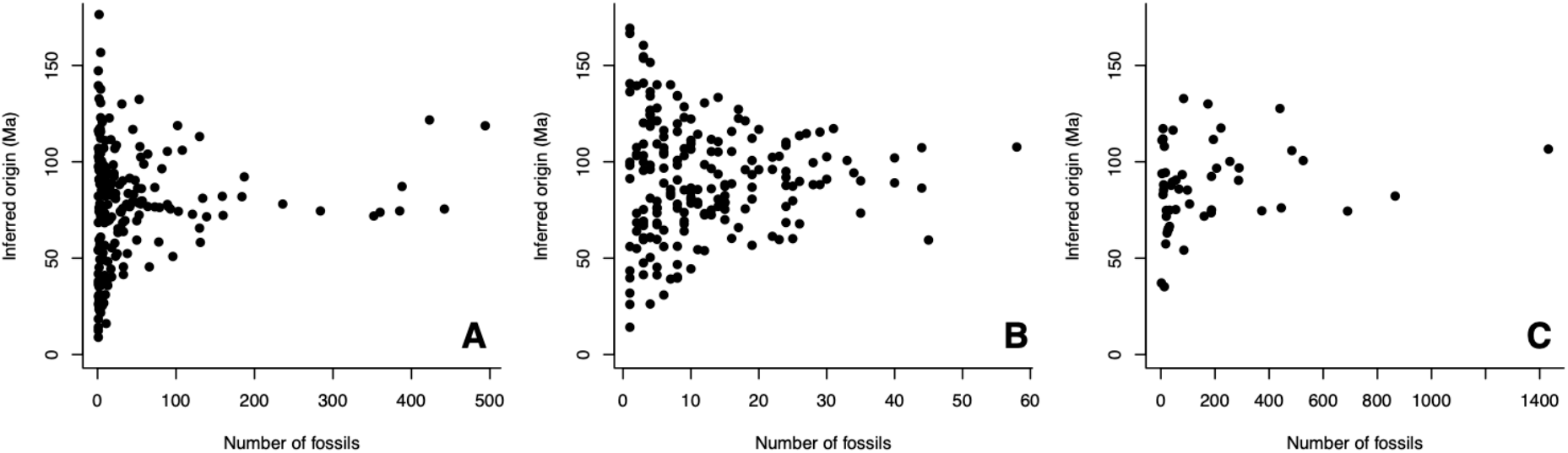
Plots of inferred mean age versus number of fossils in each family in **A** Silvestro et al.^2^, **B** Our simulated set of 198 “families” all of true age 120 Ma and **C** Our order level analysis. Note that especially in **A** and **B**, very old and very young families are characterized by few fossils, suggesting their placement there results from sampling variability (cf. Fig. 2c of Silvestro et al.^2^ which shows a similar distribution with respect to relative errors in their simulated data).

Ref. 2 reports the probability that at least one family emerged before the Cretaceous to be 0.998, by treating the age of each family as independent. Using the same procedure, our simulated results give a probability of 0.99996, despite all our clades actually originating in the Cretaceous. This nonsensical result illustrates the point that statistical treatment of all families as providing independent samples of the age of the angiosperms is incorrect. If there were no fossil data at all, with a uniform prior from 0-300 Ma on the age of the angiosperms, the probability of a pre-145 Ma origin should be 0.517, not 1-0.483^198^ (essentially 1) assuming independence. Thus, this calculation is not a useful guide to the age of crown-group angiosperms.

As Silvestro et al.^2^ recognize, their analysis is sensitive to incorrect assignment of fossil taxa to families with a sparse record, and indeed several cases are problematic. To take two from families inferred to extend into the Jurassic, they place the Early Cretaceous lobate leaf *Potomacapnos*^5^ in Papaveraceae (eudicots). However, the original description only cautiously compared it to Papaveraceae, and it shares more similarities in venation with the Early Cretaceous lobate leaf *Mesodescolea*, which appears to represent an extinct basal angiosperm line^6^. Similarly, Late Cretaceous flowers assigned to the mycoheterotrophic Triuridaceae (monocots) have smooth boat-shaped monosulcate pollen, unlike the spiny spherical inaperturate pollen of extant triurids^7^.

Altogether, Silvestro et al.^2^ infer pre-Barremian mean ages (> 131 Ma) for eight angiosperm families, but of these only Nymphaeaceae and Cabombaceae are early-divergent lineages. Tellingly, these are the only families out of the eight with > 4 fossil records, suggesting the early ages of the other six are a result of the artefacts we have discussed.

If we take the results of Silvestro et al.^2^ at face value, they imply that many eudicot lineages (united by their distinctive tricolpate pollen, modified into tricolporate and triporate within the clade) extend deep into the Jurassic. This would conflict strongly with the sequential appearance of monosulcate, tricolpate and tricolporate pollen in the Cretaceous dispersed pollen record, which represents a vastly broader stratigraphic and geographic sample of the world’s past vegetation than the record of other plant parts^1^. Finally, we note that re-running the analysis at the ordinal rather than familial level increases the number of fossils in most clades and results in a Cretaceous or younger origin for all orders (Fig. 1D).

Although basal crown-group angiosperms may well have originated in the Jurassic^1^, current evidence suggests that angiosperms did not diversify to the level implied by Silvestro et al.^2^ in that period, and the radiation of core angiosperms (mesangiosperms) may have begun in the Cretaceous. This matches the expectation^8^ that crown groups, once they emerge, diversify rapidly and should quickly enter the fossil record. We believe this provides a better solution to Darwin’s “abominable mystery” than scenarios involving an extensive period of hidden diversification.

